# Captive green iguana is a reservoir of diarrheogenic *Escherichia coli* pathotypes

**DOI:** 10.1101/624544

**Authors:** Gerardo Uriel Bautista-Trujillo, Federico Antonio Gutiérrez-Miceli, Leonel Mandujano-García, María Angela Oliva-Llaven, Carlos Ibarra-Martínez, Paula Mendoza-Nazar, Benigno Ruiz-Sesma, Carlos Tejeda-Cruz, Liset Candelaria Pérez-Vázquez, Jesús Eduardo Pérez-Batrez, Jorge E. Vidal, Javier Gutiérrez-Jiménez

## Abstract

The green iguana appears to be a reservoir for bacteria causing gastrointestinal infections in humans. The presence of diarrheogenic *E. coli* (DEC) pathotypes, however, has not been studied in this reptile. The aim of the current work was to investigate the prevalence of DEC in the intestines of 240 captive green iguanas, their phylogenetic groups, and the antibiotic susceptibility profile. *E. coli* strains were isolated from 41.7% of the intestinal content of green iguanas. DEC strains was identified in 25.9% of the screened population and were detected in the majority (62%, p=0.009] of those reptiles carrying *E. coli* strains. Among DEC strains, STEC strains carrying the *stx1* gene were the most prevalent pathotype isolated (38.7%), followed by EAEC and ETEC (27.4% each). Genetic markers of DEC strains belonging to the EHEC pathotype were not detected. More than a half of DEC strains were classified into the Clade I-II phylogroup (64.5%), followed by the phylogroup A (14.5%). The antibiotic susceptibility method demonstrated that a high proportion of DEC strains were resistance, or non-susceptible, to carbenicillin, amikacin, and ampicillin (85, 74, and 66%, respectively). We conclude that the green iguana kept in captivity is a reservoir of DEC strains bearing resistance to first-line antibiotics, including penicillins. Given the increase presence of the green iguana in Latin American households, these reptiles represent a potential source of transmission to susceptible humans and therefore a potential source of gastrointestinal disease.

**Importance:** Latin-American countries present a high burden of diarrheal disease. In this part of the world, besides common pathogens, diarrheal diseases are also caused by pathogenic types of *E. coli* referred as “diarrheagenic *E. coli*” (DEC). While inhabitants of Latin American countries suffer of self-limiting diarrhea when infected with DEC, a main target of these strains are tourists from developed countries who are not exposed to DEC strains. Efforts are in place to decrease the burden of DEC-associated diarrheal disease. As such, this study investigated a potential reservoir of DEC strains that had been underestimated, the green iguana. These reptiles are very common in Latin American households and are found virtually everywhere in Mexico, Central and South America. We found that 25% of green iguanas carried DEC strains in their intestines. We also demonstrated a high prevalence of antibiotic resistance in these strains, posing a potential risk to humans.

## Introduction

*Escherichia coli* is a commensal bacteria found in the gastrointestinal tract of human and mammals. Its ability to acquire virulence genes has originated strains that cause serious gastrointestinal infections in humans as well as animals (1). *E. coli* pathogenic strains are responsible for approximately 56 million cases of diarrhea causing 0.2 million annual deaths worldwide, most of it in children between 2 and 5 years of age (2).

Based on the virulence traits of this Gram-negative bacterium and the location of the infection within the human host, pathogenic *E. coli* strains are classified in Diarrheogenic *E. coli* (DEC) and Extra intestinal *Escherichia coli* (ExPEC). This last group includes uropathogenic *E. coli* (UPEC) and neonatal meningitis *E. coli* (NMEC). The DEC group includes Enterotoxigenic *E. coli* (ETEC), Enteropathogenic *E. coli* (EPEC), Enteroinvasive *E. coli* (EIEC), Shiga toxin producing *E. coli* (STEC), Enteroagregative *E. coli* (EAEC) and diffusely adherent *E. coli* (DAEC) (3). Among the DEC pathotypes, STEC infections cause diarrhea, and hemorrhagic colitis but the infection ocassionally progress to hemolytic uremic syndrome (HUS) that despite severe sequela can cause death (4). STEC virulence is mainly based on its ability to produce two isoforms of the Shiga toxin (Stx): Stx1 and Stx2, related to infections in human, mainly subtypes Stx2 (5). Multiple STEC serotypes have been reported in outbreaks of human disease, being the most frequent STEC O157, followed by non-O157 serotypes including O26, O103, O111, O21, O45 and O145 strains (6).

It have been reported that bovines, ovines, goats, pigs, dogs and cats are reservoirs of different DEC strains (7). Ruminants, for example, are an important reservoir of DEC strains with their feces representing a key source of contamination of water and food that cause infections to humans. Several epidemiological studies have demonstrated the zoonotic potential of STEC strains. For instance, a study conducted in Belgium analzying diarrheic calves found that 58% of STEC strains were capable of inducing the attaching/effacement (A/E) lesion, followed by EPEC strains (38%); the O26 and O111 serogroups were the most frequent (47.5 and 30%, respectively) (8). Another study found a prevalence of 0.7% of STEC serotype O103:H2 strains in feces of Norwegian sheep (9). A work conducted in Iran showed the presence of STEC strains in healthy claves and goats (26.3 and 27.5%, respectively)(5).

Recently, in Mexico, a high prevalence of STEC (40.7%) and ETEC (26.7%) was observed in dairy cows, besides reporting the presence of *E. coli* serotype O157:H7 and O104:H12 (10). The presence of DEC strains in wild animals seems to be low in comparison to their prevalence in humans and ruminants. EPEC (typical and atypical) and STEC strains have been isolated from captive wild birds in Brazil (11, 12). A study that surveyed animals from zoos in India demonstrated a low prevalence of DEC strains in wild ruminants (STEC 7.14%; EPEC 1.58%), in non-ruminant animals (STEC 3.48%; EPEC 5.81%) and wild birds (EPEC 5.84%) (13).

In Mexico and in other South-American countries, the green iguana (*Iguana iguana*) has played an important role in the economy of some regions of these Latin America countries as these animals are sell for human consumption but also kept as pets. These factors led to a dramatic decline of iguana populations the last few years. Important efforts, however, to preserve the species have emerged in Mexico such as a special protection is in place according to the Mexican Official Norm NOM-059-SEMARNAT-2010 (14) as well as efforts to promote their conservation by means of the Wildlife Management Units (WMU) (15).

Iguana species can carry in their intestines *Salmonella* species (16, 17) and *Escherichia coli* (18–20). The prevalence of *E. coli* in iguana species ranges from 40 to 70 %. For example, in the Ricord’s iguana (*Cyclura ricordi*) the prevalence reported was 50% (21), while in the land iguana (*Conolophus pallidus*) from the Galapagos Island it was 70.83 % (18). Regarding the green iguana, studies have reported an *E. coli* prevalence of 40 % (20) and 53.2 % (17). To the best of our knowledge, whether the green iguana can carry DEC pathotypes has not been previously investigated and it was the main motivation for this study.

Therefore, the goal of this study was to investigate the prevalence of DEC strains colonizing the intestines of captive green iguanas. To figure out whether a subset of DEC strains are associated to intestinal colonization of the green iguana, we also determined their phylogenetic group as well as their susceptibility to first-line antibiotics.

## Material and methods

### Green iguana population

From autumn 2015 to spring 2017, a total of 240 captive iguanas were selected from the WMU (UMAs for the acronym in Spanish) as follows: 112 specimens from the Istmo-Costa region and 128 iguanas from the Metropolitan region of Chiapas, Mexico. The reptiles were safely retrieved from their respective cages and were classified in accordance to their age: 126 juvenile iguanas (6 to 18 months of age) and 114 adults (more than 36 months of age). As juvenile iguanas present physical characteristics that do not allow distinguishing males from females, sex was determined only within the adult population. Male iguanas presented swelling of the hemipenis (females lack of this characteristic) and bigger femoral pores than females (22). The study utilized 55 females and 59 males.

### E. coli strains isolation

Intestinal content was collected in accordance to approved guidelines (23). This study was approved by the Committee for Animal Care of the University Autonomous of the Chiapas state (approval ID number 06/VET/RPR/269/16). Briefly, a sterile swab was introduced into the cloaca and after rotating gently, it was placed in Stuart agar gel (Copan) for its transportation to the laboratory. Swabs were streaked immediately onto Eosin and Methylene Blue agar (BD-BBL) and incubated at 37° C for 24 h. Lactose fermenting colonies were selected and analyzed by standard biochemical tests. *E. coli* strains were also genetically identified by PCR amplification of the *uidA* gene, that encodes for β-glucuronidase specific for *E. coli* (24).

### Diarrheogenic E. coli pathotypes, serogroups and phylogenetic group’s identification by polymerase chain reaction (PCR)

#### Reference E. coli strains

The following reference strains were used: *E. coli* strain (ATCC^®^ 25922™) as a negative control, while EAEC 042 (044:H18), ETEC H10407 (O78:H11), EPEC E2348-69 (O127:H6), STEC EDL933 (O157:H7) and EIEC E11 (O124NM) prototype strains were used as a positive control in PCR reactions. Strains belongs to the bacterial collection at the University of Sciences and Arts of the state of Chiapas (25).

#### Bacterial genes analysis

To obtain bacterial DNA, a pool of 4-5 isolated colonies were resuspended in 1 mL of deionized water and then boiled for 10 min. The bacterial lysate was centrifuged at maximun speed for five minutes and DNA-containing supernatant was separated and stored at −80°C until used. Specific gene-targets for Enteroagregative *E. coli* (EAEC) (*aap*, *agg*R and AAprobe genes) were amplified by a technique described by J. F. Cerna et al. (26). For Enterotoxigenic *E. coli* (ETEC) (*lt* and *st genes*), Enteropathogenic *E. coli* (EPEC) (*bfp*A and *eae*A genes), Shiga- toxin producing *E. coli* (STEC) (*stx*1 and *stx*2 genes) and Enteroinvasive *E. coli* (EIEC) (*ial gene*), we utilized targets and conditions described by C. López-Saucedo et al. (27). The *eae* and *hly*A genes, encoding for intimin and α-hemolysin, were amplified as established by E. Oswald et al. (28) and H. Schmidt et al. (29), respectively. The STEC strains were further analyzed in order to detect the O26, O45, O103, O111, O113, O121 and O145 antigens. To this end, primers amplifying the *wzx* gene (O-antigen flippase) in the O-antigen gene clusters were utilized as described by C. DebRoy et al. (30). The O157 antigen was investigated by amplifying the *rfb* gene (O-specific polysaccharide), according to A. W. Paton and J. C. Paton (31).

*E. coli* strains were further classified, using a quadruplex PCR method, into seven phylogroups: A, B1, B2, C, D, E, F, and the cryptic Clade I. This method is based in the presence or absence of *chuA, yjaA, arpA, trpA* genes and the TSPE4 C2 DNA fragment. PCR conditions were published by O. Clermont et al. (32) with primers shown in Table 1. PCR reactions were run in a thermal cycler C1000 (BioRad) and PCR products were analyzed through electrophoresis in agarose gel (2%) ran at 80 V for 1 h. Agarose gels were stained with SyberGreen^®^ (Invitrogen), and photographed with the Molecular Imager^®^ Gel Doc™ XR System (BioRad). The lambda molecular weight markers (100 and 1000 pb) were used (Invitrogen, USA).

**Table 1.**
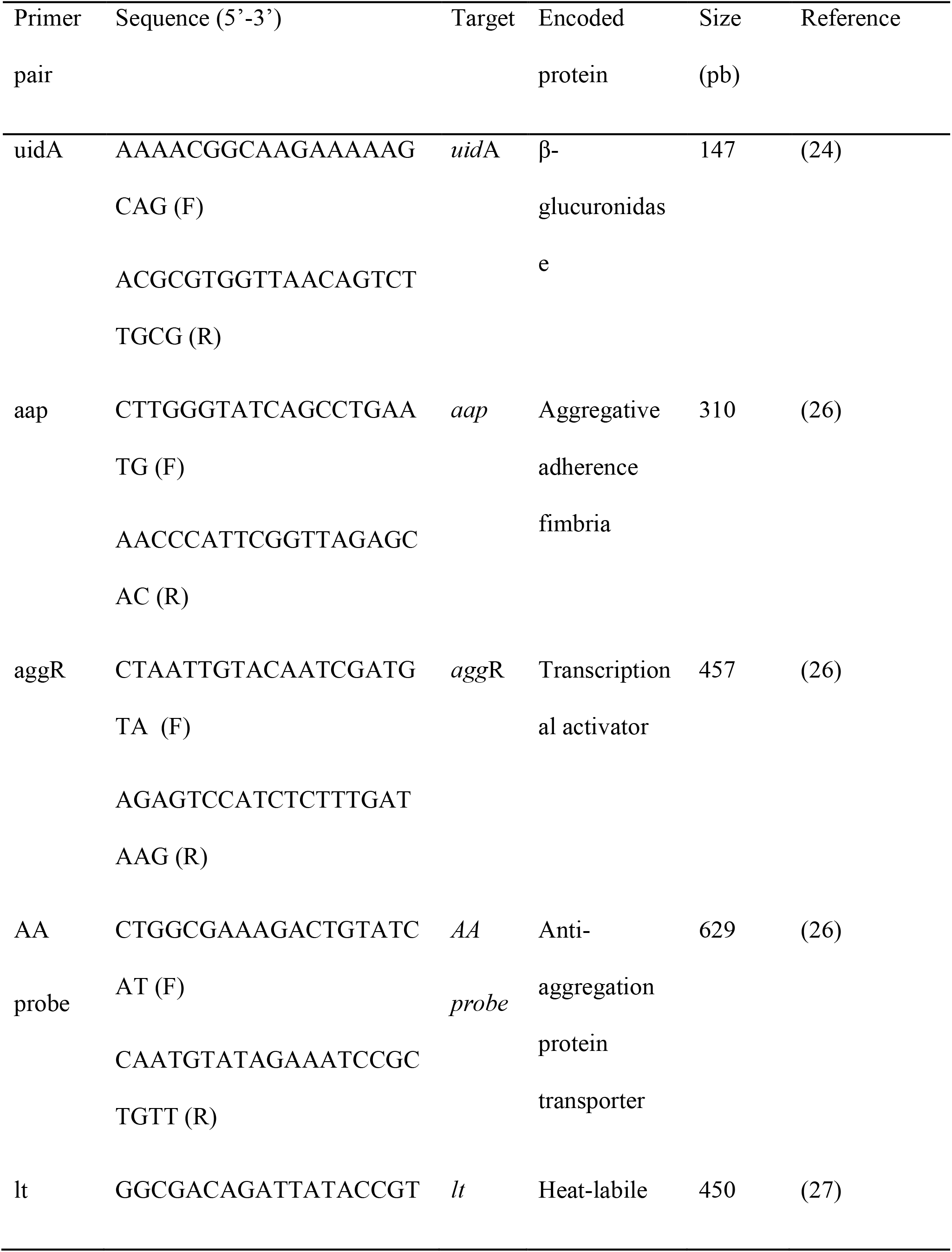
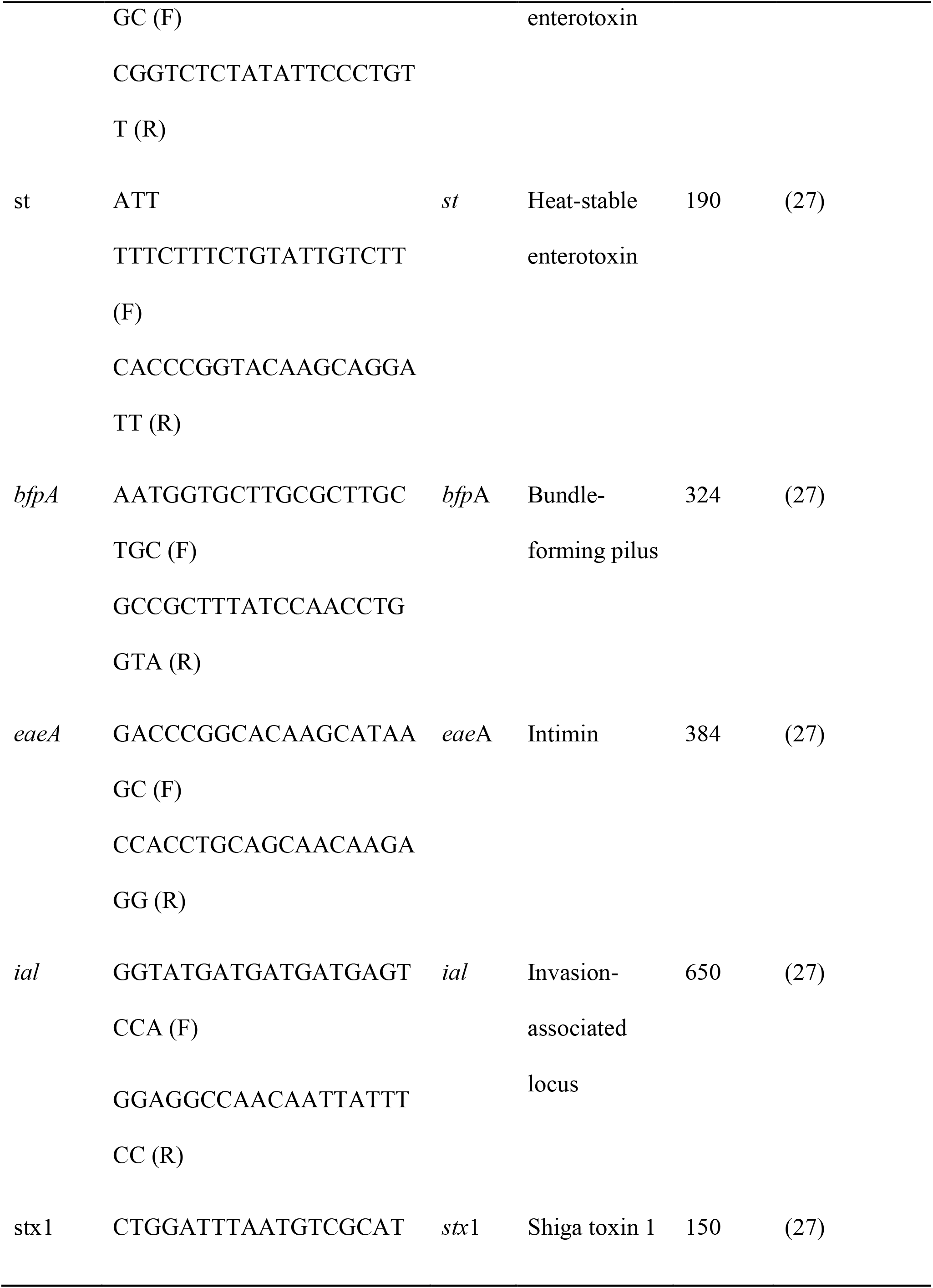
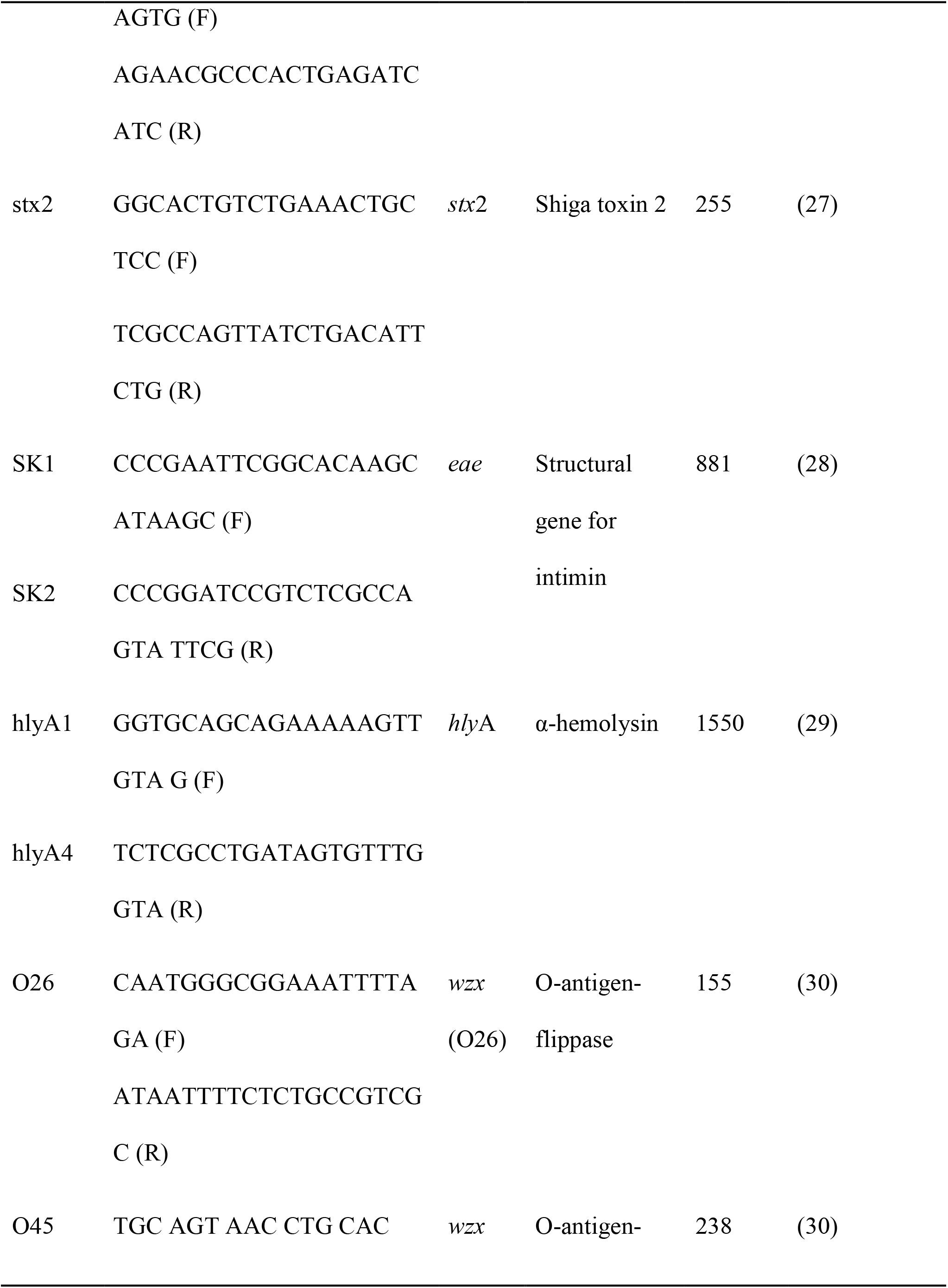
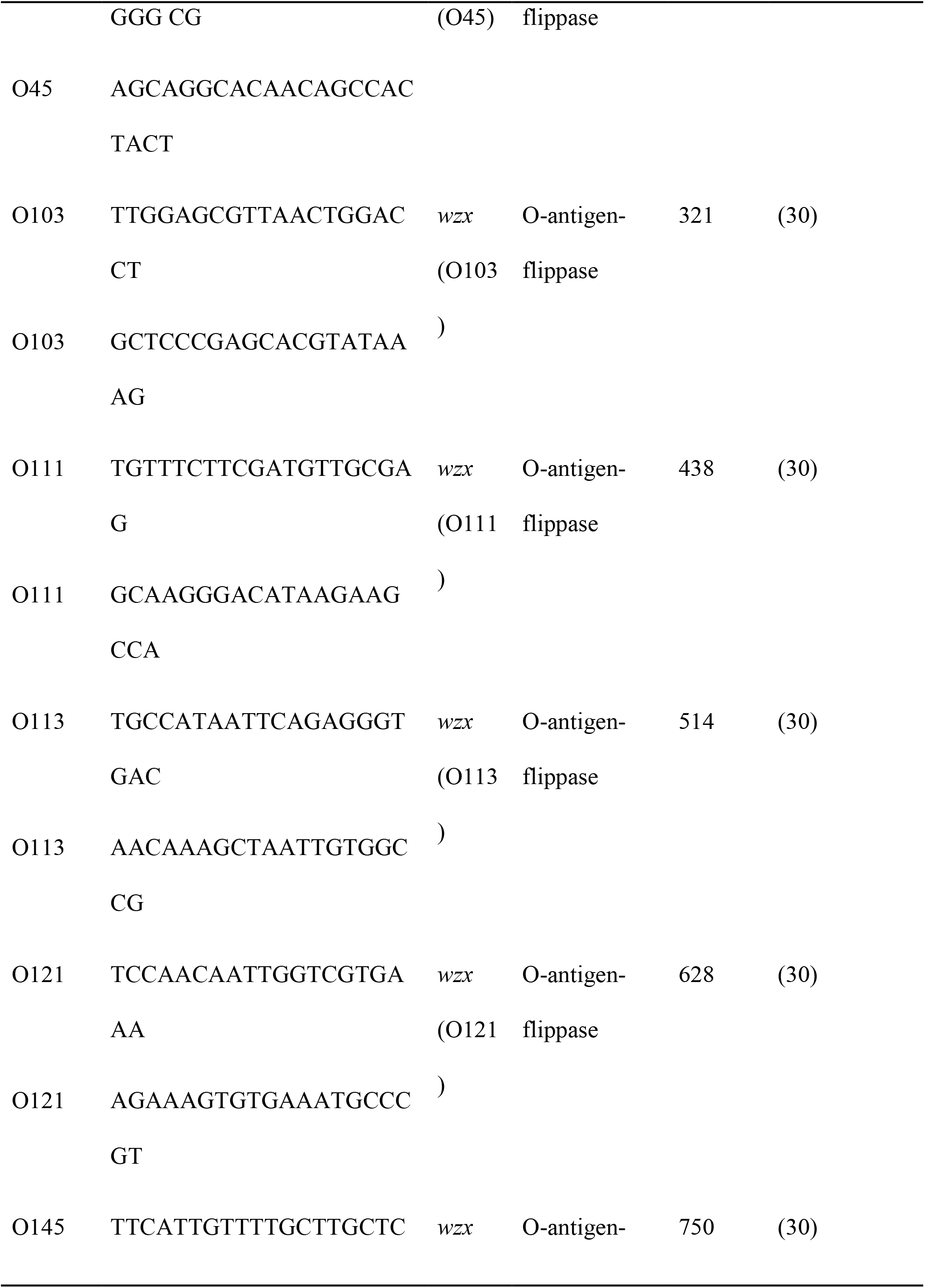
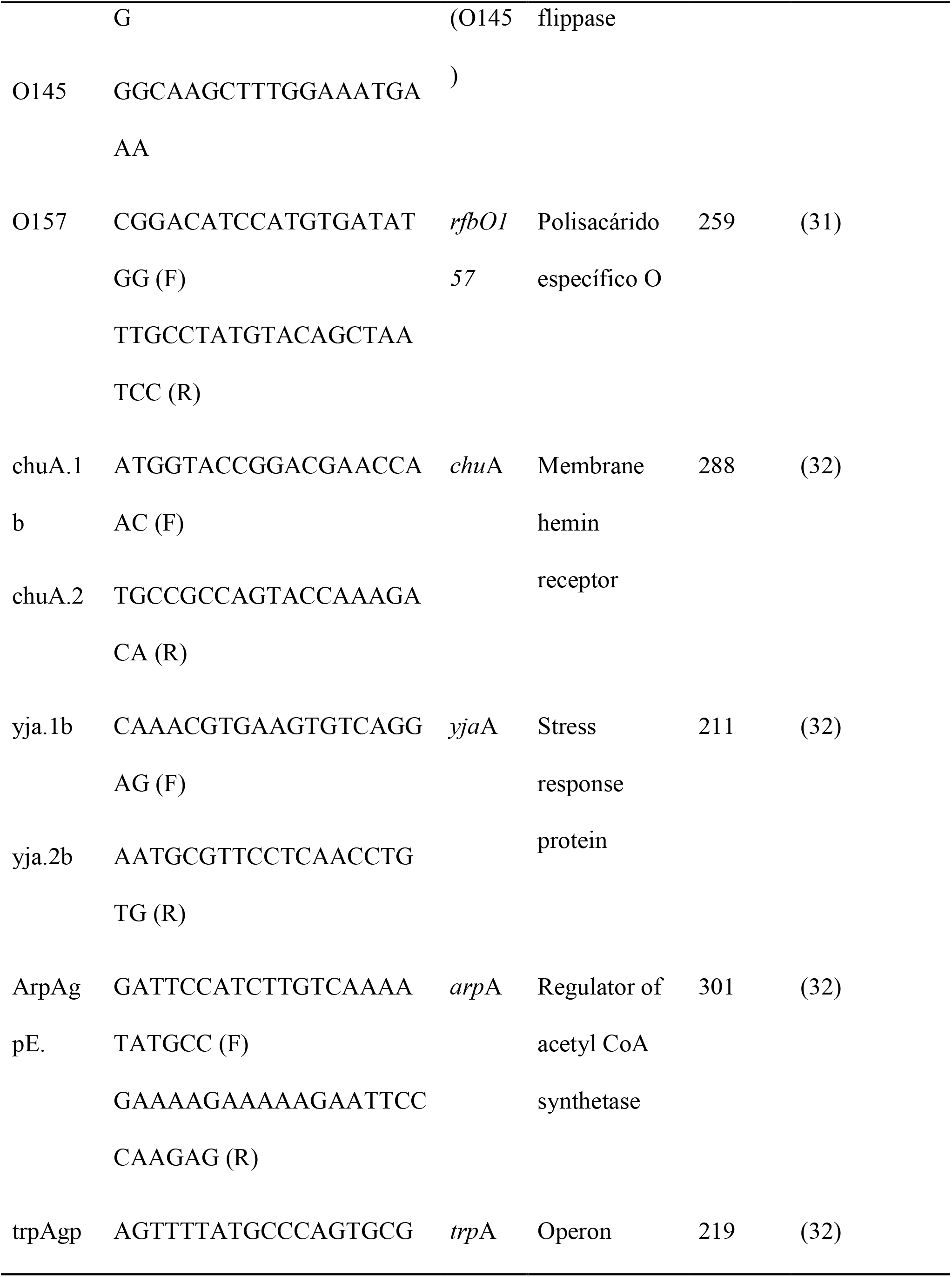
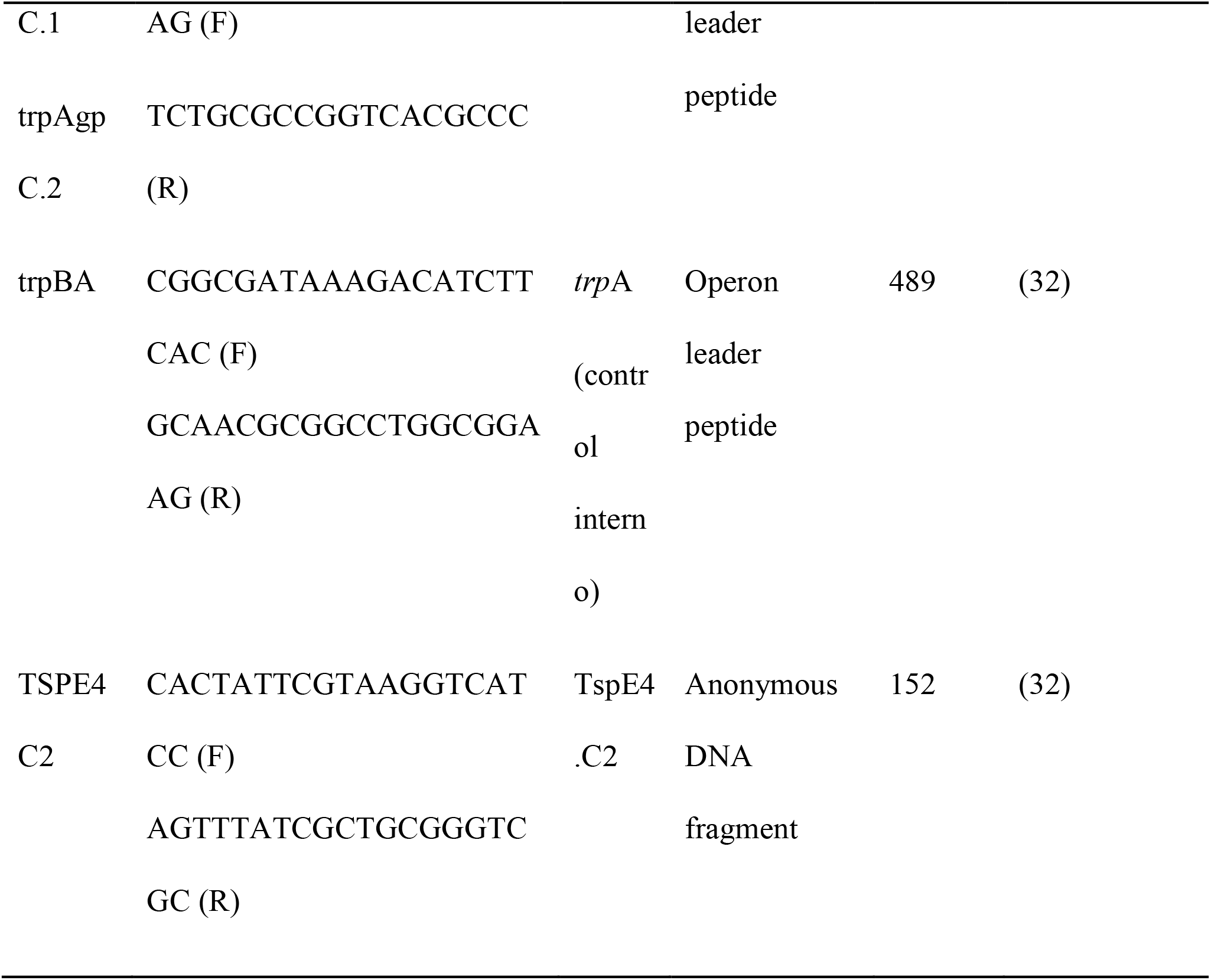
Primers used in this study.

### Antimicrobial susceptibility analysis

The disk-diffusion method was utilized following recommendations by the Clinical Laboratory Standards Institute (33). The following antimicrobial susceptibility test discs (BD BBL™ Sensi-Disc™) were utilized: amikacin (AMK;30 μg), ampicillin (AMP;10 μg), carbenicillin (CAR;100 μg), cefotaxime (CTX;30 μg), cephalotin (CEF;30 μg), chloramphenicol (CHL;30 μg), ciprofloxacin (CIP;5 μg), gentamicin (GEN;10 μg), netilmicin (NET;30 μg), nitrofurantoin (NIT;300 μg), norfloxacin (NOR;10 μg) and trimethoprim-sulfamethoxazole (SXT;25 μg). Strains resistant to β-lactams were further analyzed with amoxicillin-clavulanic acid discs (AMC;20/10 μg).

### Statistical Analysis

Prevalences of *E. coli* pathotypes, serogroups, phylogenetic groups, and susceptibility to antibiotics were analyzed by descriptive statistics. Proportions of typical molecular markers among DEC strains were analyzed by binomial test. The analysis between categorical variables was performed using the Fisher Exact test or Chi-Square test (when the frequencies were higher than 5) establishing a significance level when *p*< 0.05. All statistical analyses were conducted using the IBM SPSS statistics software (Chicago, SPSS Inc.).

## Results

### *Diarrheagenic E. coli* identification

We isolated *E. coli* strains from 41.7% (N=100) of fecal samples collected from 240 iguanas kept in captivity. DEC strains was identified in 25.9% of the screened population of green iguana and were detected in the majority [62% (N=62), *p*=0.009] of those reptiles carrying *E. coli* strains (N=100). Among DEC strains, STEC (40.3%) was the most prevalent category followed by EAEC (27.4%, with a significative frequency of the *aap* gene) and ETEC (27.4%). EPEC strains had the lowest prevalence (4.9%) (Table 2).

**Table 2.** Virulence genes identified in DEC strains isolated from *I. iguana* in Chiapas, Mexico.

| DEC category | Virulence factor | % (N) | *p* * |
| --- | --- | --- | --- |
| STEC |  | 40.3 (25) |  |
|  | *stx1* | 38.7 (24) | 0.000002 |
|  | *stx2* | 1.6 (1) |  |
| EAEC |  | 27.4 (17) |  |
|  | *aap* | 27.4 (17) | 0.0001 |
|  | AA probe | 1.6 (1) |  |
| ETEC |  | 27.4 (17) |  |
|  | *lt* | 19.3 (12) | 0.359 |
|  | *st* | 11.3 (7) |  |
| EPEC |  | 4.9 (3) |  |
|  | *bfpA* | 4.9 (3) | 1.000 |
|  | *eaeA* | 3.2 (2) |  |
| Total |  | 100.0 (62) |  |

508 \*Compares differences within each DEC category for carrying a virulence marker.

Our PCR approach revealed a statistically significant higher frequency of STEC strains carrying the *stx*1 gene in comparison to those carrying *stx*2. The *eae* and *hly*A virulence genes were not amplified in any of these STEC strains suggesting that they did not belong to the EHEC category. Accordingly, O-antigen genes carried by EHEC’s most common serotypes (e.g., O26, O45, O103, O111, O113, O121, O145 and O157) were not amplified in any of these strains.

### Investigation of phylogenetic groups

Given that these iguanas are in captivity we shought to investigate whether DEC strains were clonal by obtaining their phylogenetic group. Among these *E. coli* strains (N=62), the clade I or II phylogroup was the most predominant (64.5%), followed by phylogroups A and B2 accounting for the 14.5 and 3.2% of strains, respectively. Some of DEC strains (17.8%) could not be assigned to a known phylogroup (Table 3). Thus, DEC isolated from green iguanas appear to be clonal since the majority of strains were classified in clade I or II.

**Table 3.** Phylogenetic groups of *Escherichia coli* strains isolated in captive green iguana in Chiapas.

| Phylogenetic<br>groups | *E. coli* strains % (N) |  |  |  |  | *p* * |
| --- | --- | --- | --- | --- | --- | --- |
|  | STEC stx1 | EAEC | ETEC | EPEC | STEC stx2 |  |
| A | 25.0 (6) | 0.0 (0) | 17.6 (3) | 0.0 (0) | 0.0 (0) | 0.156 |
| B2 | 0.0 (0) | 5.8 (1) | 5.9 (1) | 0.0 (0) | 0.0 (0) | 0.392 |
| Clade I-II | 75 (18) | 70.5 (12) | 41.2 (7) | 66.6 (2) | 100.0 (1) | 0.114 |
| Unknown | 0.0 (0) | 5.8 (4) | 35.3 (6) | 33.3 (1) | 0.0 (0) | 0.004 |

\*Comparing associations between phylogenetic groups and DEC's category.

There was not association between the prevalence of *E. coli* pathotypes and the age or sex of the iguanas. A similar trend was observed among *E. coli* pathotypes and phylogroups. Regarding the geographic region were these reptiles are kept in captivity, a statistically significant association was observed between STEC strains isolated from iguanas kept at the metropolitan (i.e., urban) region in comparison to those from the coast (i.e., rural) (Table 4).

**Table 4.**
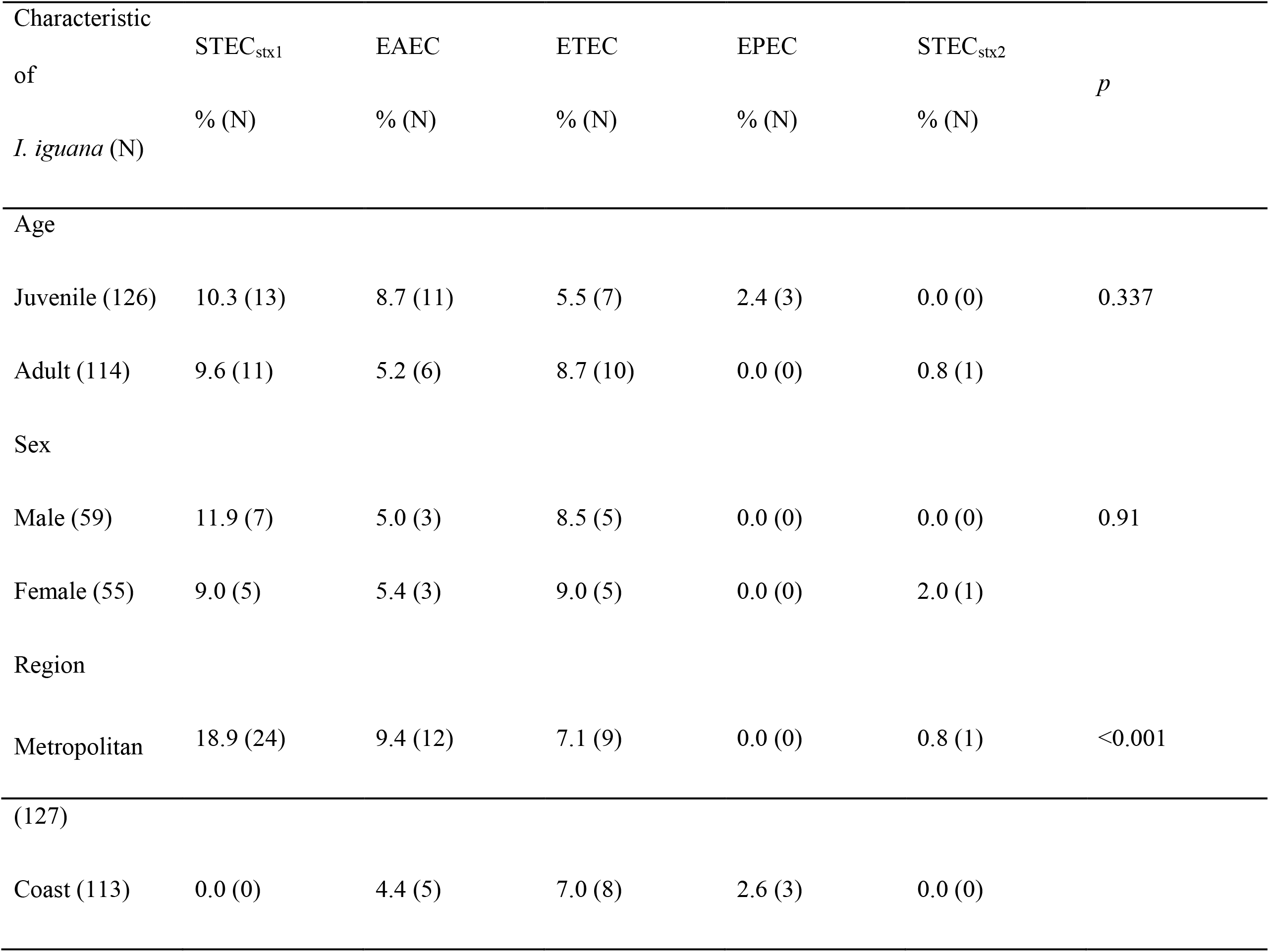
Associations among *E. coli* strains and some characteristics of the green iguana in Chiapas.

### Antimicrobial susceptibility of DEC strains

Because these DEC strains represent a potential source of contamination for humans and therefore a source of gastrointestinal disease, we investigated their susceptibility to antibiotics utilized to treat gastrointestinal disease caused by Gram negative bacteria. More than a half of DEC strains were resistant to carbenicillin (85%, resistant and intermediate), amikacin (74%, resistant and intermediate), and ampicillin (66%, resistant and intermediate) (Fig. 1). All strains non-susceptible to ampicillin were susceptible to amoxicillin-clavulanic acid (data not shown).

**Figure 1.**
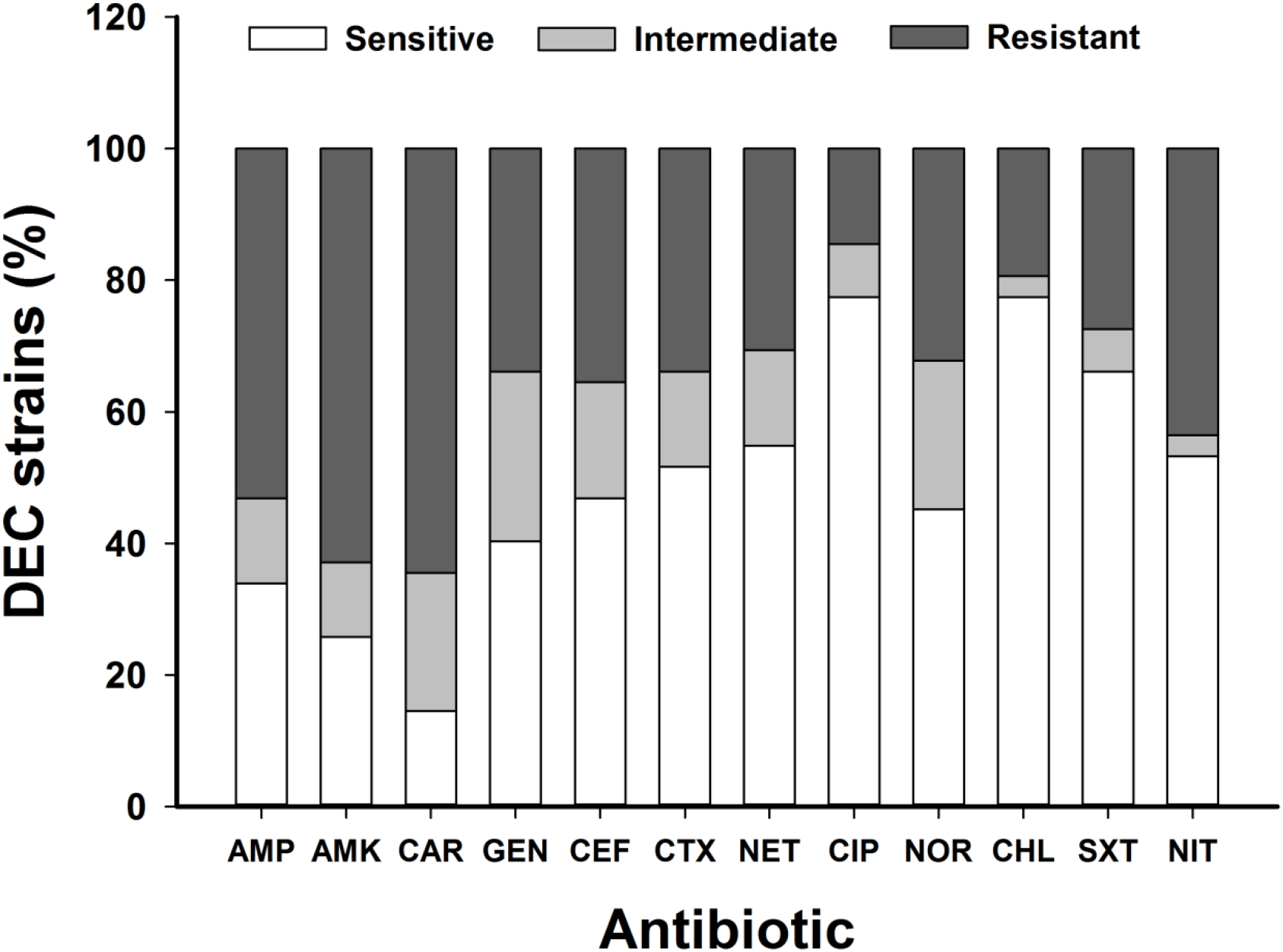
Antibiotic susceptibility patterns of DEC strains (N=62) isolated from *I. iguana* in Chiapas, Mexico. AMP; Ampicillin, AMK; Amikacin, CAR; Carbenicillin, GEN; Gentamicin, NET; Netilmicin, CEF; Cephalotin, CTX; Cefotaxime, CIP; Ciprofloxacin, NOR; Norfloxacin, CHL; Chloramphenicol, SXT; Trimethoprim-sulfamethoxazole, NIT; Nitrofurantoin.

## Discussion

*Escherichia coli* is an ubiquitous Gram negative bacteria considered as one of the main constituent of the gastrointestinal tract of animals (34), humans (35), as well as reptiles such as turtles (36) and snakes (37). This trend is also observed in species of iguana, as it is documented in several studies including a work by W. Sylvester et al. (20) who reported a prevalence of 40% of *E. coli* strains in green iguana from Granada, West Indies. A similar prevalence (50%) of this bacterium was found in Ricord’s iguana (*Cyclura ricordii*) from Enriquillo Lake of the Isla Cabritos National Park, (Dominican Republic) (21). These results are comparable to the prevalence reported in this work (41.7%). Another group that studied the Land iguana (*Conolophus pallidus*) living in the Galapagos Island revealed a high prevalence of this Gram negative rod (91.7 %) (18).

Despite *E. coli* strains are harmless to their mammal host, certain strains collectively known as DEC strains exhibit an ability to cause disease in humans and animals by virtue of their virulence factors, allowing the replication, dissemination, and damage to susceptible hosts (3, 38). Nevertheless, there is almost no information about the presence and prevalence of *E. coli* pathotypes, such as DECs in the intestines of reptiles (39). This study revealed that the majority of *E. coli* strains isolated from *I. iguana* carried genes encoding virulence factors associated to diarrhea in humans (62%) such as those carry by STEC, EAEC, or ETEC pathotypes. Acquisition of these DEC strains can be a result of coprophagy, which is a behavior seen in iguanas that it is required for biochemical digestion of the plant cells. Several studies have demonstrated that *E. coli* pathotypes are isolated from dairy cows (7), sheep (40) and pigs (41). To the best of our knowledge, however, the presence of DEC strains in the intestine of iguanas had not been reported. In Mexico, high prevalence of STEC (40.7%) and ETEC (26.7%) strains in bovine’s feces from the states of Jalisco, Sinaloa, and Sonora (10) have been reported. Contamination with feces of these mammals represent a vehicle for *E. coli* infections in human, whom may acquire these strains through the consumption of food or water contaminated with feces of animals by direct contact with infected animals specially those utilized as pets (42).

Captivity in urban settings along with the mentioned coprophagy should account for the presence of DEC strains in the intestine of iguanas (43). These observations are supported by the fact that a statistical significant association was observed in this study between intestinal carriage of STEC strains and iguanas kept in captivity at a metropolitan region of Chiapas. Reptiles may be in closer contact with human feces because of the crowdedness of an urban setting. Our findings agree with those by W. Sylvester et al. (20) whom demonstrated an association between the prevalence of *E. coli* carriage and certain geographic regions. We hypothesize that factors such as the weather, season of the year (summer and autumn), as well as a potential hygiene deficiency when handling iguanas (faecal contamination in water and food in cages) favors the acquisition of DEC bacteria (44, 45). We found similar carriage prevalences of the pathogenic variants of DEC strains in *I. iguana* according to sex and age.

In the current study, the prevalence of STEC carrying the *stx1* gene was higher than that of those carrying the stx2 gene (Table 1). None of these strains carried the *eae* and *hlyA* genes, neither genes that encode for the following O antigens: O26, O45, O103, O111, O113, O121, O145 and O157. Stx toxins represent the key virulence factors for diarrheal disease caused by STEC strains (5). Stx2 toxin is more virulent than Stx1 because that toxin is responsible for causing Hemolytic Uremic Syndrome in humans (46). Similar to the current work, a study conducted in Isla Cabritos National Park (Dominican Republic) and Granada (West Indies) failed to isolate EHEC O157:H7 from *I. iguana* (20, 47). EHEC strains belonging to serotypes O157, 026, O145 and O111 have been isolated from cases of HUS in humans from USA and Europe (48, 49). Therefore, our data indicate that the green iguana kept in captivity is a reservoir for Stx1-producing STEC strains but not for EHEC strains.

The present study also revealed that the *E. coli* isolates were grouped mainly in Clade I-II. As such, our results agree with studies demonstrating that environmental *E. coli* strains belong to different cryptic clades (50). To our knowledge, there are no studies about the phylogenetic classification of *E. coli* isolated from reptiles by using the current quadruplex phylogrouping method (32).

Regarding susceptibility to antbiotics, >50% of DEC strains were non-susceptible to six out of 12 antibiotics assessed incuding a high resistance to penicillins, aminoglycosides and some of the broad-spectrum antibiotics cephalotin and norfloxacin (Figure 1). These data are worrisome since treatment failure is expected in the event of a zoonosis of gastrointestinal illness. While DEC strains had not been isolated from iguanas until this study, in other studies isolating normal flora *E. coli* from iguanas captured at a Caribbean island, including that one by W. Sylvester et al. (20), authors reported susceptibility to different antibiotics in *E. coli* isolates. The non-susceptibility to antibiotics observed in DEC strains isolated from iguanas in Chiapas indirectly indicates that resistance is a consequence of the acquisition of strains from humans since these reptiles are not treated with antibiotics during their life-span. Moreover, the resistance phenomenon is not common between bacteria isolated from wildlife animals (18). Then, these findings suggest a microbiological impact caused by the conditions of captivity.

The green iguana is utilized as a pet in some Latin American countries, including in South Mexico. Animals used with this purpouse are considered as a potential source of zoonosis and, as observed in this study, they may also bear strains with resistance to a commonly used antibiotics that may complicate infections or even outbreaks of zoonotic origin.

## Conclusions

The current study demonstrated the presence of diarrheagenic *E. coli* (mainly STEC) in the green iguana kept in captivity. The findings of the current work might be extremely useful for the sanitary efforts that are made during the green iguana handling, with the purpose of reducing its potential as a reservoir for pathogenic bacteria such as DEC strains. Additional studies about the genetic lineage of the pathotypes found in the green iguana will allow us to understand the clonal origin and predict possible bacteria outbreaks in reptiles kept in captivity, which may represent a risk for public health.

## Acknowledgements

We want to thank Dayanf Fernanda Saldaña Molina, Eglantina Corzo Cobos, Paola Gómez Velázquez, Brenda Ovando Diaz, Gabriel Nucamendi Moguel and Jessica Estrada Vázquez, for their contribution in the data collection. We also thanks the assistance provided by personnel of the Wildlife Management Units of the State of Chiapas, Mexico. Financial support was partially obtained by the Program for Strengthening Educational Quality of the University of Science and Arts of the State of Chiapas (grant number C/PFCE-2016-07MSU0002G-13-43).

